# The heterogeneous functional architecture of the posteromedial cortex is associated with selective functional connectivity differences in Alzheimer’s disease

**DOI:** 10.1101/700856

**Authors:** Wasim Khan, Ali Amad, Vincent Giampietro, Emilio Werden, Sara De Simoni, Jonathan O’Muircheartaigh, Eric Westman, Owen O’Daly, Steve C.R. Williams, Amy Brodtmann

## Abstract

The posteromedial cortex (PMC) is a key region involved in the development and progression of Alzheimer’s disease (AD). Previous studies have demonstrated a heterogenous functional architecture of the region, with different subdivisions reflecting distinct connectivity profiles. However, little is understood about PMC functional connectivity and its differential vulnerability to AD pathogenesis. Using a data-driven approach, we applied a constrained independent component analysis (ICA) on healthy adults from the Human Connectome Project (HCP) to characterise the distinct functional subdivisions and unique functional-anatomic connectivity patterns of the PMC. These connectivity profiles were subsequently quantified in the Alzheimer’s Disease Neuroimaging Initiative (ADNI) study, to examine functional connectivity differences in (1) AD patients and cognitively normal (CN) participants and (2) the entire AD pathological spectrum, ranging from CN participants and participants with subjective memory complaints (SMC), through to those with mild cognitive impairment (MCI), and finally, patients diagnosed with AD. Our findings revealed decreased functional connectivity in the anterior precuneus, dorsal posterior cingulate cortex, and the central precuneus in AD patients compared to CN participants. Functional abnormalities in these subdivisions were also related to high amyloid burden and lower hippocampal volumes. Across the entire AD spectrum, functional connectivity of the central precuneus was associated with disease progression and specific deficits in memory and executive function. These findings provide new evidence showing that specific vulnerabilities in PMC functional connectivity are associated with large-scale network disruptions in AD and that these patterns may be useful for elucidating potential biomarkers for measuring disease progression in future work.

## Introduction

Alzheimer’s disease (AD) is a progressive neurodegenerative disorder characterised by a decline in memory and cognitive functions. The disease is related to the pathological accumulation of aggregated amyloid depositions and hyperphosphorylation of structural proteins which lead to metabolic alterations, functional loss, and structural changes in the brain. Convergent evidence across neuroscience disciplines suggests that proteinopathies progress trans-synaptically along brain networks, with neuronal dysfunction topographically spreading from a region of focal onset to non-adjacent regions in a predictable pattern manifesting over several years (Greicius & Kimmel, 2012; Liu et al., 2012). The location and distribution of pathogenic processes such as the accumulation of amyloid deposits has been consistently mapped to a network of heteromodal regions collectively known as the default mode network (DMN) (Buckner et al., 2005).

Some of the earliest and consistent pathological changes observed in AD are evident in the posteromedial cortex (PMC) – an integrated hub region important for episodic memory encoding and retrieval (Vannini et al., 2013). Large decrements in glucose metabolism of the PMC and a vulnerability to amyloid pathology are consistent early features of AD which are known to manifest prior to the onset of clinical symptoms (Minoshima et al., 1997; Mintun et al., 2006). In the later stages of AD pathogenesis, a disruption to connections between the PMC and large-scale memory and visual networks can also be observed (H. Y. Zhang et al., 2009). This suggests that an accurate characterisation of PMC function is of vital importance to the understanding of AD development and progression. However, little is known about the pattern of PMC functional connectivity with distributed large-scale brain networks across different stages of the AD pathological spectrum.

Previous anatomical studies have demonstrated that the PMC consists of highly diverse cytoarchitectonics and is functionally heterogenous (Margulies et al., 2009). Yet, the PMC is often treated as having a homogenous functional architecture in studies of the DMN, despite its diverse patterns of functional connectivity (Y. Zhang et al., 2014). Recent work characterising the functional architecture of the PMC has shown that it consists of several different functional subdivisions that are associated with multiple large-scale brain networks at rest (Leech, Braga, & Sharp, 2012; Margulies et al., 2009; S. Zhang & Li, 2012). In particular, the posterior cingulate cortex (PCC), which lies in the medial part of the inferior parietal lobe, exhibits distinct cytoarchitectonics with functional separation into dorsal and ventral areas (Leech, Kamourieh, Beckmann, & Sharp, 2011; Vogt, Vogt, & Laureys, 2006). This dorsal region of the PCC demonstrates strong connectivity with the DMN and other large-scale networks, including the frontoparietal network involved in executive control and the salience network for attention, thus implicating its role in modulating global network metastability (Hellyer, Scott, Shanahan, Sharp, & Leech, 2015; Leech et al., 2012). In contrast, the ventral region of the PCC is highly integrated within the DMN, particularly with key medial prefrontal and temporal nodes and is understood to be involved in internally directed cognition, such as memory retrieval and planning (Dastjerdi et al., 2011; Leech et al., 2012). As a result, studies have suggested that the PMC plays a central associative role across a wide-spectrum of integrated functions with evidence of its involvement in sensorimotor processing, cognitive functioning, and the processing of visual information (Hutchison, Culham, Flanagan, Everling, & Gallivan, 2015). However, only a handful of studies have examined the different functional subdivisions of the PMC in AD (Cauda et al., 2010; Xia et al., 2014). Furthermore, no studies have addressed, to the best of our knowledge the susceptibility of these subdivisions across the entire AD spectrum as well as the relationship between PMC subdivisions and other well-established disease markers of AD pathology.

A few resting-state fMRI (rsfMRI) studies in AD have parcellated the PMC to examine its intrinsic functional architecture, however most have used *a priori* defined cortical seed regions to characterise its complex functional neuroanatomy (Cauda et al., 2010; Dillen et al., 2016; Margulies et al., 2009; Wu et al., 2016). Recent work has highlighted the advantages of multivariate data analysis methods for a finer and more detailed parcellation of brain regions (De Simoni et al., 2018; Leech et al., 2012).

Here, we used high-resolution rsfMRI data from HCP to fractionate the PMC into its subdivisions using a constrained independent component analysis (ICA) method and characterise its unique patterns of functional connectivity with large-scale brain networks. Subsequently, these detailed maps of the PMC were used to compare functional connectivity differences in AD using the publicly available and widely phenotyped Alzheimer’s Disease Neuroimaging Initiative (ADNI) study. Specifically, the aim of this study was to examine functional connectivity changes in the different subdivisions of the PMC across the AD spectrum (*N*=155), ranging from cognitively normal (CN) participants and participants with subjective memory complaints (SMC) through to those with mild cognitive impairment (MCI), and AD. We first characterised the functional connectivity of the PMC by examining disease-specific changes in AD patients compared to CN participants. The functional connectivity of PMC subdivisions that were disrupted in AD were subsequently tested for their association with amyloid burden and hippocampal volume measurements. Furthermore, we hypothesised that brain networks of PMC subdivisions that were strongly implicated in cognition would be associated with specific deficits in memory and executive function.

## Materials and Methods

For this study, data was obtained from HCP (Van Essen et al., 2012) (http://www.humanconnectome.org/) for defining PMC subdivisions and their functional connectivity patterns in healthy unaffected young adults. These detailed functional maps and results were later used to assess functional connectivity patterns in AD patients and CN participants from the ADNI database (http://adni.loni.usc.edu/). For the purposes of simplicity, we will describe these different datasets in the same order as the workflow from our analysis pipeline.

### Human Connectome Project

#### Data and preprocessing

rsfMRI data for 100 randomly selected healthy adult participants (Age range: 22-36 years; 46 males) were obtained from the HCP S1200 data release. Participants had no documented history of mental illness, neurological disorder, or physical illness with known impact upon brain functioning. This cohort of participants were selected such that there were no related participants within the cohort due to concerns over the heritability of neural features (Glahn et al., 2010). Data were acquired on a Siemens Skyra 3T scanner housed at Washington University in St. Louis (TR = 720 ms, TE = 33.1 ms, spatial resolution = 2 ×2 × 2 mm^3^), collected in four separate 15-min runs on two different days (two per day). Each rsfMRI run consisted of 1,200 volumes which totalled to 4,800 volumes (over the four runs). Quality assurance and quality control procedures of HCP for rsfMRI data have been described previously (Marcus et al., 2013). The data we obtained had been minimally preprocessed by HCP (Glasser et al., 2013), as well as denoised to remove non-neural spatiotemporal components (Griffanti et al., 2014; Salimi-Khorshidi et al., 2014) and high-pass filtered (Satterthwaite et al., 2013).

#### Defining posteromedial cortex functional subdivisions

A constrained ICA was performed using the minimally preprocessed HCP datasets in the FMRIB Software Library (FSL) (5.0.11) using the MELODIC tool (Beckmann, DeLuca, Devlin, & Smith, 2005). A temporal concatenation group ICA was constrained to extract components within a PMC mask, defined *a priori* using the Harvard-Oxford probabilistic atlas. The PMC was constructed by selecting PCC and precuneus regions with a voxel probability threshold greater than 20% signal intensity. rsfMRI data within the mask of the PMC were spatially smoothed using an 8mm full width at half maximum Gaussian (FWHM) kernel following masking. In order to determine the ideal number of ICA components to decompose the PMC into its distinct functional subdivisions, we performed a reproducibility analysis to assess the procedures trade-off between granularity and noise using the mICA toolbox (Moher Alsady, Blessing, & Beissner, 2016). Reproducibility analyses were performed for an ICA dimensionality range of 2-20 ICA components and showed that 10 ICA components were best representative of the underlying HCP data. Consequently, MELODIC was run to extract 10 ICA components for the PMC. Further details regarding the reproducibility analysis are provided in the Supplementary Methods section.

#### Characterising cortical functional connectivity of posteromedial cortex subdivisions

A variant of the dual regression procedure in FSL (Beckmann, Mackay, Filippini, & Smith, 2009; Filippini et al., 2009) was used to characterise the cortical functional connectivity patterns of PMC subdivisions derived from ICA (Fig 1A). The final statistical maps from this approach provide a whole-brain measure of functional connectivity between each voxel in the brain and the decomposed ICA signal of the PMC. Further details of this approach are described in previous work (De Simoni et al., 2018; Leech et al., 2012, 2011). To define cortical functional connectivity patterns of each PMC subdivision in the HCP dataset, a general-linear model was applied in two steps. Firstly, ICA spatial maps of the PMC subdivisions were regressed against whole-brain rsfMRI data, resulting in subject-specific time-courses for each ICA spatial map (i.e. each PMC subdivision). This step served to generate a subject specific timecourse for each spatial map of the ICA whilst controlling for the variance explained by the other spatial maps. Secondly, whole-brain rsfMRI data were modelled in another general-linear model using the previously calculated time-courses as regressors. This resulted in subject-specific whole-brain spatial maps of network coherence for each PMC subdivision (Fig 1B). Group average maps were calculated using a general linear model. To account for multiple comparisons, the TFCE method was used (Smith & Nichols, 2009) with 5000 permutations.

**Figure 1.**
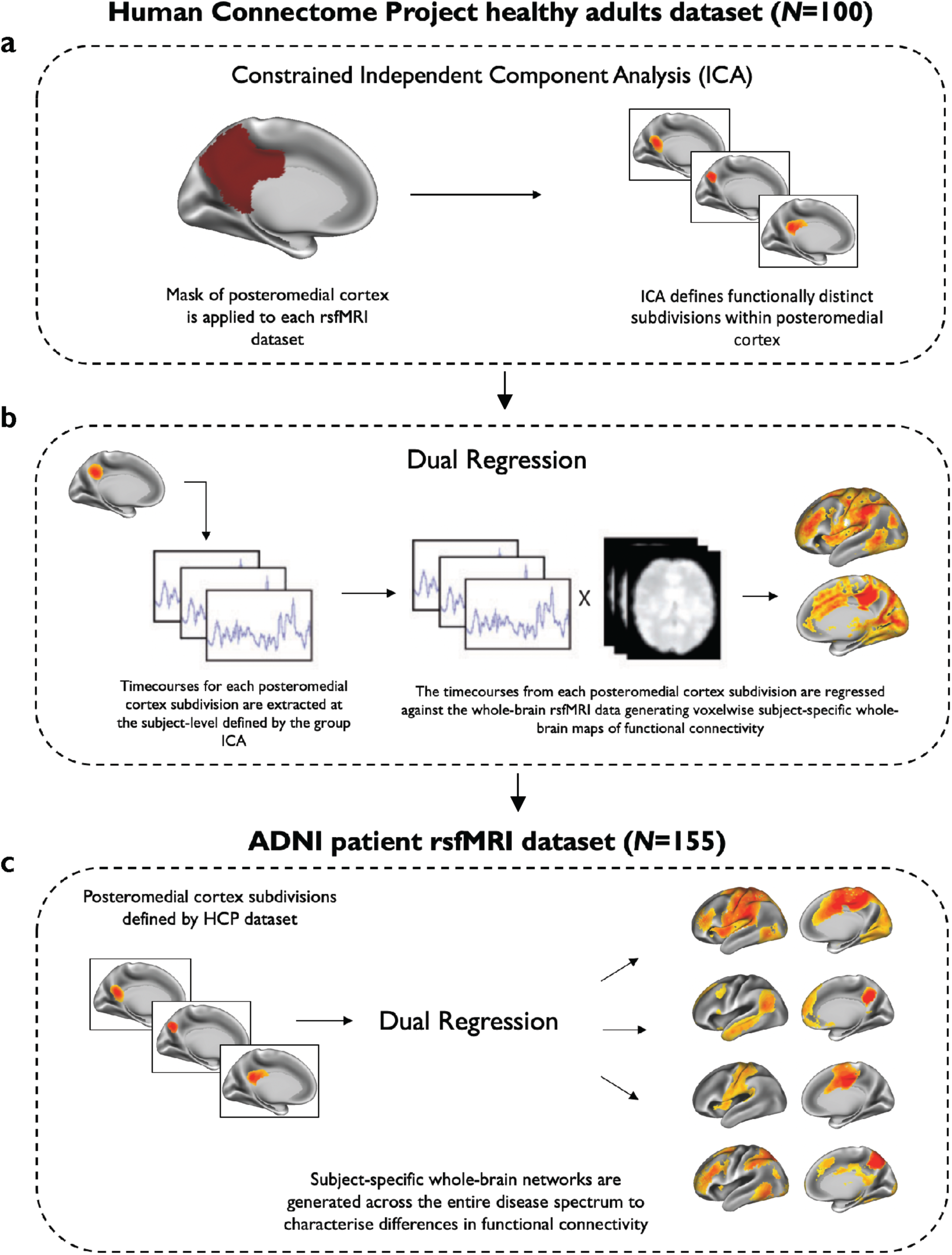
A schematic representation of the major steps involved in the functional connectivity analysis of the posteromedial cortex. **(a)** An ICA approach was used to fractionate subdivisions of the posteromedial cortex (PMC) by constraining the analysis to voxels within a pre-defined PMC mask in the HCP dataset. **(b)** In the same HCP dataset, a dual-regression analysis was used to define the functional connectivity patterns of each PMC subdivision by correlating its activity with voxels in the rest of the brain. **(c)** PMC subdivisions defined in the HCP dataset were used to characterise whole-brain functional connectivity of brain networks identified in the ADNI patient dataset (*N*=155). This dataset included cognitively normal (CN) participants, participants with subjective memory complaints (SMC), early mild cognitive impairment (MCI) participants, late MCI participants, and patients with Alzheimer’s disease (AD). The brain networks shown here in **(c)** were constrained to voxels of the same brain networks identified in the HCP dataset, shown in **(b)**.

### Alzheimer’s Disease Neuroimaging Initiative patient dataset

Data used for this study was obtained from the ADNI database. ADNI is a multi-centre longitudinal biomarker study that has enrolled over 1,500 CN participants, people with early or late stages of MCI, and patients with early AD (www.adni-info.org). ADNI was approved by the institutional review board and ethics committees of participating institutions, and written informed consent was obtained according to the Declaration of Helsinki from all participants or their next of kin.

#### Participants

All participants were downloaded from the ADNI-2 database (June 2018). Baseline scans were identified from all images that had undergone quality control implemented by the Mayo Clinic (*N*=827) (Jack et al., 2015). We selected all first available scans for each participant as their baseline scan (*N*=225). Based on our inclusion criteria, several participants were excluded (*N*=53) to ensure that we only retained data of a reasonable quality. This included any scans with motion parameters exceeding 1.5mm of translation and/or rotation (*N*=11), scans with an image quality rating > 3 (image quality rated as:: 1=excellent; 2=good; 3=fair; 4=poor) (*N*=36), presence of microhaemorrhages or cysts (*N*=3), or any other uncertainties in diagnosis (*N*=3). Of the remaining 172 participants, if amyloid imaging or *APOE* genotyping was unavailable (*N*=17), these participants were not included for further analysis. PET amyloid imaging was performed using Florbetapir. A measure of amyloid burden was calculated from frontal, cingulate, parietal, and temporal regions and was averaged and divided by a whole cerebellum reference region to create a standardised uptake value ratio. A threshold of 1.11 was used to define amyloid positivity. This has been described in detail previously (Landau et al., 2013). Hippocampal volumes were calculated using FreeSurfer (version 6.0) (http://surfer.nmr.mgh.harvard.edu/) (Fischl, 2012).

A full description of the structural and functional image preprocessing steps for the ADNI study are provided in the Supplementary Methods section. The final dataset of participants used in this study is summarised in Table 1. A complete list of participant scans used in this study is provided in Supplementary Table 1.

**Table 1.** Demographic characteristics and metadata of the ADNI rsfMRI dataset (*N*=155).

| | CN<br>($N=34$) | SMC<br>($N=24$) | EMCI<br>($N=43$) | LMCI<br>($N=31$) | AD<br>($N=23$) | *P*-Value |
| --- | --- | --- | --- | --- | --- | --- |
| Baseline Age (years) | $75.3 \pm 6.3$ | $71.9 \pm 5.3$ | $71.1 \pm 6.9$ | $71.2 \pm 7.7$ | $72.9 \pm 7.7$ | 0.062 |
| Sex (Male %) | 14 (41) | 10 (42) | 17 (40) | 20 (65) | 12 (52) | 0.20 |
| Years of Education | $16.1 \pm 2.0$ | $16.7 \pm 2.9$ | $15.8 \pm 2.8$ | $16.6 \pm 2.6$ | $15.6 \pm 2.7$ | 0.28 |
| *APOE* $\epsilon 4$ carriers (%) | 11 (32) | 8 (33) | 22 (51) | 14 (45) | 18 (78) | **0.01** |
| CDR sum-of-boxes | $0.06 \pm 0.2$ | $0.04 \pm 0.1$ | $1.4 \pm 1.0$ | $1.7 \pm 1.0$ | $4.5 \pm 1.2$ | **< 0.001** |
| MMSE | $28.9 \pm 1.1$ | $29.1 \pm 0.9$ | $28.3 \pm 1.7$ | $27.5 \pm 1.5$ | $22.3 \pm 2.5$ | **< 0.001** |
| ADAS-Cog11 | $5.6 \pm 2.5$ | $5.6 \pm 2.3$ | $7.8 \pm 3.3$ | $10.9 \pm 4.1$ | $24.3 \pm 7.8$ | **< 0.001** |
| RAVLT forgetting (%) | $39.1 \pm 24.4$ | $37.7 \pm 22.4$ | $54.7 \pm 29.1$ | $67.6 \pm 25.9$ | $95.5 \pm 10.4$ | **< 0.001** |
| Trail Making Test B | $89 \pm 64$ | $80 \pm 42$ | $100 \pm 48$ | $112 \pm 65$ | $209 \pm 86$ | **< 0.001** |
| Amyloid Florbetapir SUVR | $1.15 \pm 0.20$ | $1.13 \pm 0.18$ | $1.21 \pm 0.21$ | $1.26 \pm 0.25$ | $1.45 \pm 0.18$ | **< 0.001** |
| Hippocampal volume (mL) | $7.6 \pm 0.84$ | $7.7 \pm 1.1$ | $7.4 \pm 0.9$ | $7.2 \pm 1.3$ | $6.1 \pm 1.1$ | **< 0.001** |
| Framewise displacement | $0.15 \pm 0.09$ | $0.17 \pm 0.09$ | $0.14 \pm 0.07$ | $0.13 \pm 0.05$ | $0.13 \pm 0.06$ | 0.66 |

Results are displayed as mean ± sd. A Kruskal-Wallis rank sum test was used for comparison of group differences in continuous variables. Categorical variables were inspected for group differences using a Fisher's exact test with *P*-values generated using 2000 Monte Carlo simulations.
*Abbreviations:* AD = Alzheimer's disease; ADAS-cog11 = 11-item Alzheimer's Disease Assessment Scale-Cognitive subscale; CDR = Clinical Dementia Rating scale sum-of-boxes; CN = Cognitively Normal; EMCI = Early Mild Cognitive Impairment; LMCI = Late Mild Cognitive Impairment; MMSE = Mini-Mental-State Examination; RAVLT = Rey Auditory Verbal Learning Test; SUVR = Standardised Uptake Value Ratio.

#### Assessment of memory performance and executive functioning

As a measure of memory performance, we used Mini-Mental-State Examination (MMSE) scores and the 11-item Alzheimer’s Disease Assessment Scale-cognitive subscale (ADAS-Cog11) scores. Neuropsychological measures of verbal memory and executive function included Rey Auditory Verbal Learning Test (RAVLT) percentage forgetting scores and Trail Making Test B scores, respectively.

#### Examining posteromedial cortex functional connectivity patterns in AD patients and healthy controls

The 10 ICA spatial maps of the PMC subdivisions defined in the HCP dataset were used to characterise whole-brain PMC functional connectivity patterns in the ADNI dataset (*N*=155). This was performed using the same dual regression procedure described above. Whole-brain maps of PMC functional connectivity were constrained to voxels identified within the same corresponding whole-brain map from the HCP cohort (Fig 1C). These functional maps were subsequently transformed back to their original native space. This resulted in functional maps that contained voxelwise information about the spatial location and magnitude of functional connectivity at the individual subject level (Filippini et al., 2009; Jones et al., 2016). The average beta of these spatial maps across all voxels was extracted as a final measure of functional connectivity. To obviate potential biases resulting from differences in grey matter volume, only voxels with >0.5 probability of containing grey matter were considered. This approach was preferred over voxelwise statistical comparisons to avoid the potentially huge multiple comparison penalty associated with comparing several PMC networks across multiple groups. Summary metrics of these brain networks also provided the opportunity to extensively investigate their relationship with disease markers of AD pathology.

#### Statistical analysis

All subsequent analyses were performed using the R statistical software environment (http://www.r-project.org) (version 3.5.1) (The R Core Team, 2018). Conditions for meeting normality assumptions were tested using QQ-plots, the distribution of residuals and the absence of multicollinearity. A Kruskal-Wallis rank sum test was used to compare continuous demographic and cognitive measures between different groups and categorical variables were compared using a Fisher’s exact test. PMC brain networks that were not normally distributed were transformed using a *Z*-family of distributions (Chou, Polansky, & Mason, 1998; Johnson, 1949), after which no deviation from a normal distribution was observed via a Shapiro-Wilk normality test.

PMC functional connectivity differences were assessed between AD patients and healthy controls using a multivariate linear regression. Two different models were constructed: the first using age, sex, years of education and framewise displacement as covariates and the second additionally correcting for *APOE* ε4 genotype. We performed these multivariate regression models using a permutation testing approach in the lmPerm package (Wheeler & Torchiano, 2016) which samples all possible permutations to estimate *P*-values. Statistical significance of regression models was assessed using type II multivariate analysis of variance (MANOVA) tests and the Pillai test statistic. Next, we tested the association of PMC brain networks found to be disrupted in AD with amyloid burden and hippocampal volume. Semi-constrained step-down regression models were used with predictors including age, sex, years of education, and *APOE* ε4 genotype. For comparisons with hippocampal volume, intracranial volume measurements were always included as a predictor in the final model. Model fits were assessed using the Akaike and Bayesian Information Criterion to report the most parsimonious model.

To investigate the effect of PMC functional connectivity on cognition, we applied linear regression models on all participants at different stages of the AD pathological spectrum (*N* = 155; Table 1). Firstly, we tested the relationship between specific PMC brain networks that were found to be abnormal in AD with memory performance (i.e. MMSE) and clinical measures of disease severity (i.e. ADAS-Cog11). In additional models, we also compared disrupted PMC brain networks with executive functioning, using RAVLT percentage forgetting scores, and verbal memory measured using Trail Making Test B scores. Regression models were constructed in a similar fashion as described above.

All analyses were corrected for multiple comparisons using the Bonferroni adjustment. Only results that survived multiple comparison correction with an alpha threshold of α = 0.05 were reported in this study.

## Results

### ADNI Demographics

A total of 155 participants were selected for this study, consisting of CN participants (*N*=34), participants with SMC (*N*=24), early MCI participants (*N*=43), late MCI participants (*N*=31), and patients with AD (*N*=23) (Table 1). *APOE* ε4 carriers were significantly greater in proportion in AD patients, followed by early and late MCI participants (*P* = 0.01). Overall, AD patients and MCI participants demonstrated greater functional and cognitive impairments compared to CN participants on several clinical and neuropsychological tests (*P* = < 0.001). No significant differences were observed for sex distribution, years of education, and maximum framewise displacement of rsfMRI scans. We also did not observe any significant correlations between framewise displacement and PMC functional connectivity.

Results are displayed as mean ± sd. A Kruskal-Wallis rank sum test was used for comparison of group differences in continuous variables. Categorical variables were inspected for group differences using a Fisher’s exact test with *P*-values generated using 2000 Monte Carlo simulations. *Abbreviations*: AD = Alzheimer’s disease; ADAS-cog11 = 11-item Alzheimer’s Disease Assessment Scale-Cognitive subscale; CDR = Clinical Dementia Rating scale sum-of-boxes; CN = Cognitively Normal; EMCI = Early Mild Cognitive Impairment; LMCI = Late Mild Cognitive Impairment; MMSE = Mini-Mental-State Examination; RAVLT = Rey Auditory Verbal Learning Test; SUVR = Standardised Uptake Value Ratio.

### Posteromedial cortex fractionation reveals distinct functional subdivisions

The ICA analysis from the HCP cohort identified 10 functionally distinct and spatially overlapping subdivisions of the PMC. These subdivisions can be observed on a functional parcellation map of the PMC shown in Fig. 2a. Each PMC subdivision is also illustrated separately in Fig. 2b. To ensure that our PMC fractionation was representative of the underlying HCP data and not an artefact of the number of components chosen for ICA decomposition, we performed a reproducibility analysis to select the ideal number of ICA components for decomposing the PMC signal. Results of this analysis demonstrated that 10 ICA components were ideal for fractionating the PMC (see supplementary figure 1).

**Figure 2.**
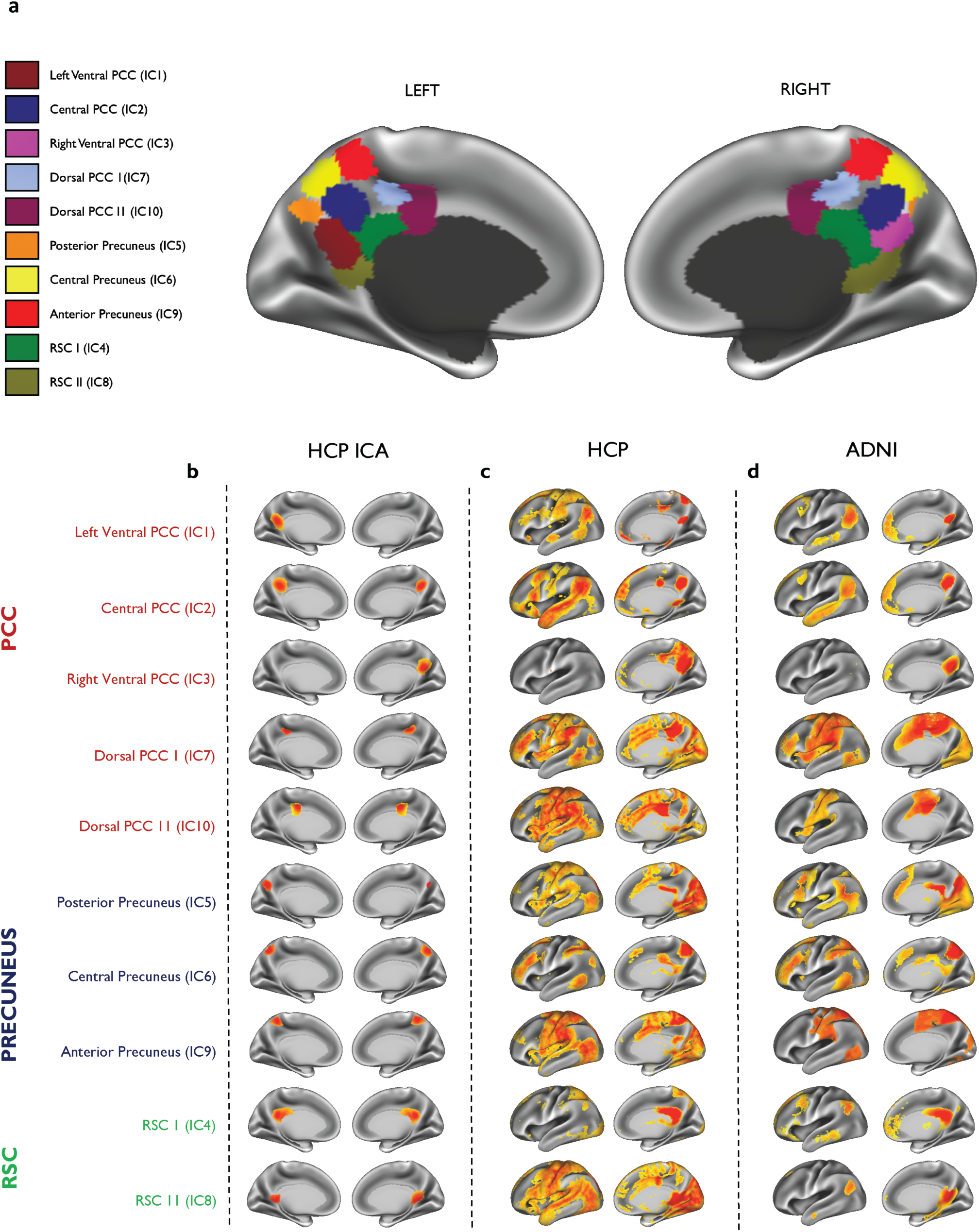
Subdivisions of the posteromedial cortex and their associated brain networks in the HCP cohort and ADNI patient dataset. **(a)** A parcellation map showing the location of all subdivisions defined in the posteromedial cortex (PMC) using the HCP dataset. **(b)** The location of each PMC subdivision is shown separately on the far left. Each subdivision (*displayed as left and right medial hemispheres*) is numbered by its ICA component and highlighted as anatomically representing the posterior cingulate cortex (PCC) in *red*, precuneus in *blue* and the retrosplenial cortex (RSC) in *green*. The whole-brain network (*displayed as left lateral and right medial hemisphere*) of the corresponding PMC subdivision is shown for **(c)** the HCP dataset and **(d)** the ADNI patient dataset (*N*=155). Warmer colours indicate areas of high functional connectivity. All maps are thresholded at *P* < 0.05 and are family-wise-error corrected for multiple comparisons.

As illustrated in Fig. 2a and 2b, five subdivisions were found to be located in the PCC, primarily in the ventral, dorsal, and anterior-dorsal parts. Three subdivisions were located in the anterior, central and posterior parts of the precuneus. Two subdivisions were found to be located in the RSC. Although PMC subdivisions shared a relatively low spatial similarity overall (*r* = 0.06-0.113), the highest spatial overlap was found between the dorsal and ventral parts of the PMC (*r* = 0.113; Fig 3a). This suggests that, despite some spatial overlap, the ICA results produced maps of considerable granularity and spatial separation.

**Figure 3.**
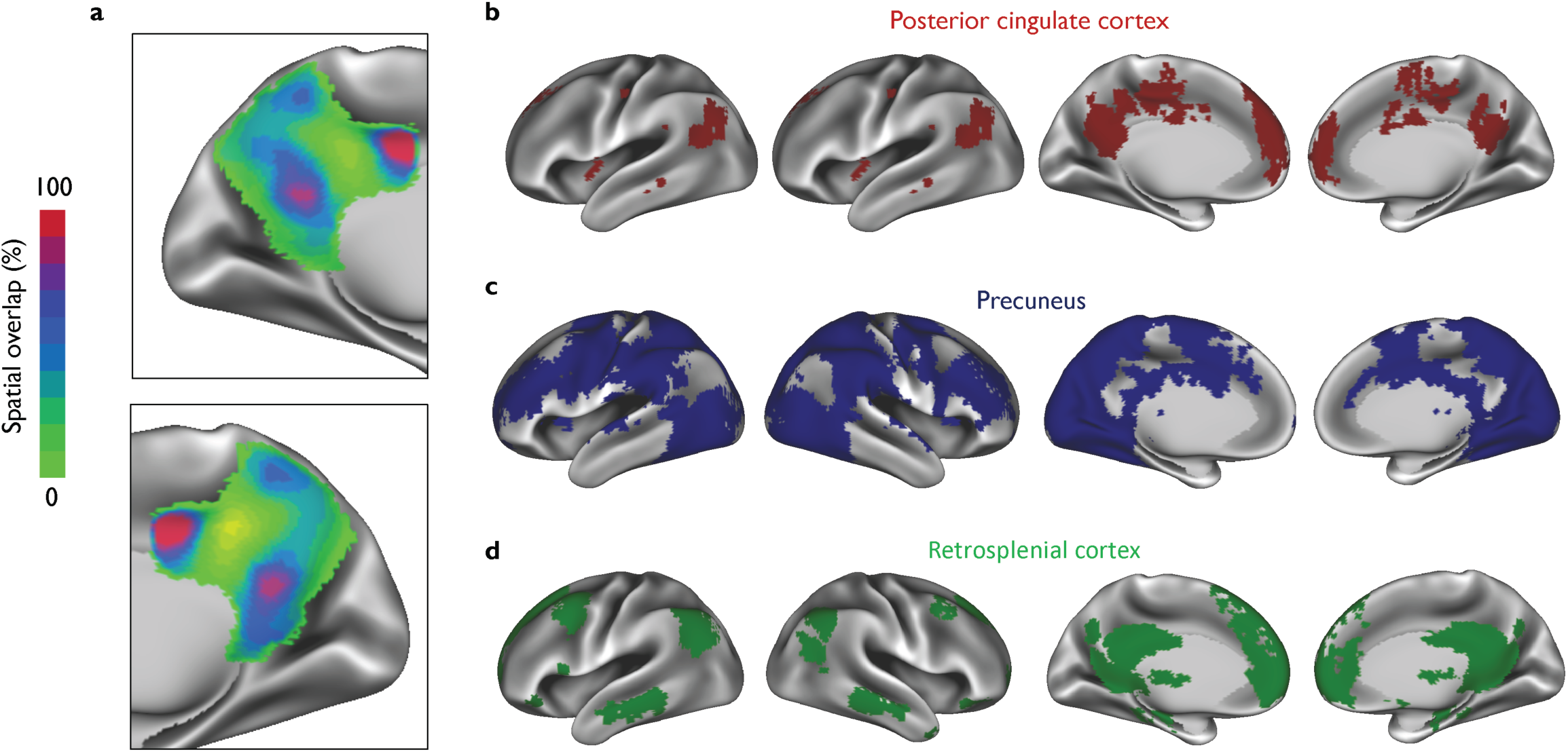
A spatial overlay map of posteromedial cortex functional subdivisions and its corresponding brain networks. **(a)** The overlap between all 10 subdivisions of the posteromedial cortex (PMC) is shown demonstrating the greatest overlap in the dorsal region of the posterior cingulate cortex (PCC), the ventral PCC and parts of the retrosplenial cortex (RSC). Also shown are spatial overlay masks of all brain networks originating from **(b)** the PCC, **(c)** the precuneus, and **(d)** the RSC. Maps are displayed as left and right lateral and medial hemispheres.

### Functional subdivisions of the posteromedial cortex are connected to several distinct brain networks

Next, we investigated whether signals from the different PMC subdivisions correlated with activity in the rest of the brain. Subdivisions of the PMC and their associated brain networks are shown in Figure 2. Our findings showed a complex functional heterogeneity of the PMC where different subdivisions were found to be associated with several distinct brain networks (Fig. 2c and Fig. 2d). The three subdivisions (IC5, IC6 and IC9) located within the precuneus revealed a distributed pattern of functional connectivity and included areas such as the frontal pole, supramarginal gyrus, temporal gyrus, occipital cortex, and occipital fusiform gyrus. For the five subdivisions of the PCC (IC1, IC2, IC3, IC7 and IC10), we found a more organised pattern of functional connectivity. Functional networks associated with IC1, IC2 and IC3 of the PCC were more DMN-like in appearance, whereas IC3 and IC7 resembled salience and frontoparietal networks covering parts of the inferior parietal regions, dorsolateral prefrontal cortex and pre-supplementary motor area. For the RSC, the two subdivisions (IC4 and IC8) showed similar patterns of functional connectivity.

We further generated spatial overlay masks of all the brain networks arising from the precuneus (Fig. 3b), the PCC (Fig. 3c), and the RSC (Fig. 3d). These were not used to compare functional connectivity differences in our study, but rather to show areas of the brain that were functionally common within anatomical regions of the PMC. Overlay masks of the precuneus reveal that more areas of the brain are functionally correlated with its different functional subdivisions. For the precuneus and RSC, overlay masks appear to be topographically similar to the DMN.

### Functional connectivity of the posteromedial cortex is reduced in Alzheimer’s disease

We compared the functional connectivity differences in PMC activity between AD patients (*N*=23) and CN participants (*N*=34). Results are described from two models, one corrected for age, sex, years of education and framewise displacement and the second additionally correcting for *APOE* ε4 genotype. These results are shown in Table 2. Comparisons were performed for the functional connectivity of all PMC subdivisions as no *a priori* hypotheses about specific network changes were postulated. The overall multivariate regression model was statistically significant (Pillai test statistic = 0.32; *P* = 0.002) and remained significant when additionally correcting for *APOE* ε4 genotype.

**Table 2.** PMC functional connectivity differences in AD patients and CN participants.

|  | *t* | Cohen's *d* | *P*-Value † | *P*-Value ‡ |
| --- | --- | --- | --- | --- |
| <u>*PCC:*</u> |  |  |  |  |
| ***Left ventral PCC (IC 1)*** | -1.83 | -0.49 | 0.07 | 0.05 |
| *Central PCC (IC 2)* | -1.76 | -0.48 | 0.08 | 0.07 |
| *Right ventral PCC (IC 3)* | 0.13 | 0.04 | 0.90 | 0.99 |
| ***Dorsal PCC I (IC 7)*** | -2.93 | -0.79 | 0.004 | 0.003 |
| *Dorsal PCC II (IC 10)* | -1.06 | -0.29 | 0.29 | 0.36 |
| <u>*Precuneus:*</u> |  |  |  |  |
| *Posterior Precuneus (IC 5)* | -1.45 | -0.39 | 0.15 | 0.35 |
| ***Central Precuneus (IC 6)*** | -3.20 | -0.86 | <0.001 | <0.001 |
| ***Anterior Precuneus (IC 9)*** | -2.68 | -0.72 | 0.008 | 0.019 |
| <u>*RSC:*</u> |  |  |  |  |
| *RSC I (IC 4)* | -0.68 | -0.18 | 0.50 | 0.69 |
| *RSC II (IC 8)* | 0.92 | 0.25 | 0.36 | 0.13 |

† Corrected for age, sex, years of education, and framewise displacement (*MANOVA Pillai test statistic* = 0.321 ; *p* = 0.002).
‡ Corrected for age, sex, years of education, *APOE* ε4 genotype, and framewise displacement (*MANOVA Pillai test statistic* = 0.329 ; *p* = 0.002).
Results have been corrected for multiple comparisons using the Bonferroni method.
Abbreviations: AD = Alzheimer's disease; CN = cognitively normal participants; IC = independent component; PCC = posterior cingulate cortex; PMC = posteromedial cortex; RSC = retrosplenial cortex

Functional networks associated with two PCC subdivisions demonstrated reduced functional connectivity patterns compared to CN participants. This included the brain network of the left ventral PCC (IC3) (*t* = −1.83; Cohen’s *d* = −0.49; *P* = 0.05) and the dorsal PCC (*t* = −2.93; Cohen’s *d* = −0.79; *P* = 0.003). For the precuneus, functional networks of the central precuneus subdivision (*t* = −3.20; Cohen’s *d* = −0.86; *P* = < 0.001) and the anterior precuneus subdivision (*t* = −2.68; Cohen’s *d* = −0.72; *P* = 0.019) were significantly reduced in AD patients. No functional connectivity differences were observed for subdivisions of the RSC. We also did not find any significant increases in functional connectivity. Linear regression analyses were performed using these brain networks that were disrupted in AD patients to determine their association with amyloid burden and hippocampal volume. Results revealed that decreased functional connectivity of the dorsal PCC, and central precuneus were associated with greater levels of amyloid burden and lower hippocampal volumes (Fig. 4).

**Figure 4.**
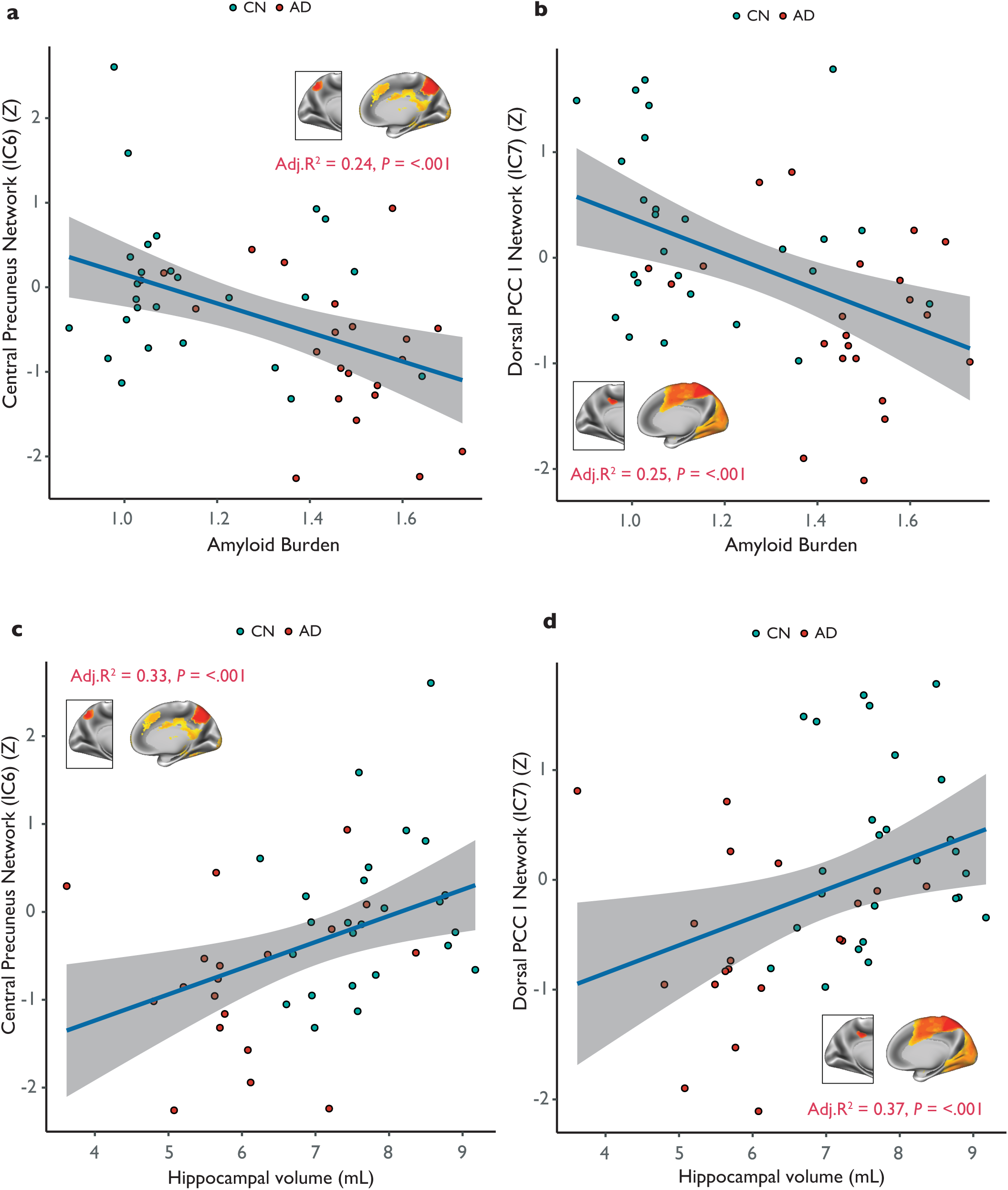
The relationship of the central precuneus and dorsal PCC networks with amyloid burden and hippocampal volume. Results are displayed for AD patients (*N*=23) and CN participants (*N*=34) in the ADNI dataset. Functional connectivity of **(a)** the central precuneus network, and **(b)** the dorsal PCC plotted versus PET Florbetapir measures of amyloid burden. Similar plots are illustrated for **(c)** the central precuneus network, and **(d)** the dorsal PCC network against FreeSurfer derived measures of hippocampal volume. The variance explained by each of the models (Adj. R-squared) and *P*-values are displayed inset. Individual data points, regression lines and 95% CI’s (grey bands) are displayed for each plot. Covariates considered in regression models included age, gender, years of education and *APOE* ε4 genotype. Models of hippocampal volume were corrected for intracranial volume measurements. AD = Alzheimer’s disease; CN = cognitively normal; PCC = posterior cingulate cortex.

### Central precuneus functional connectivity is associated with clinical disease progression, memory deficits and executive dysfunction

The central precuneus was strongly decreased in AD patients and has been previously implicated to play an integrated cognitive/associative role in the brain (Margulies et al., 2009). Therefore, we sought to determine whether aberrant functional connectivity of this PMC subdivision would be associated with disease severity and specific deficits in memory. Since no prior hypotheses were established regarding the different pathophysiological profiles of AD, we did not stratify participants on the basis of amyloid status (i.e. amyloid positive vs. amyloid negative). Instead, participants across the entire disease spectrum were included (*N*=155; Table 1), ranging from CN participants and participants with SMC, through to participants with early and late stages of MCI, and finally patients diagnosed with AD.

Results showed that weaker functional connectivity of the central precuneus was significantly associated with increasing disease severity measured using ADAS-cog11 scores (Adj. R^2^ = 0.22; *P* = < 0.001; Fig. 5b). Weaker functional connectivity of the central precuneus was also significantly associated with lower MMSE scores (Adj. R^2^ = 0.21; *P* = < 0.001; Fig. 5c). A similar association was also observed with a measure of executive function, where lower central precuneus connectivity was significantly related to higher Trail Making Test B scores (Fig. 5d). A linear reduction in central precuneus functional connectivity was also associated with a linear increase in RAVLT percentage forgetting scores, suggesting that abnormal connectivity in the central precuneus network is also associated with verbal memory deficits (Fig. 5e).

**Figure 5.**
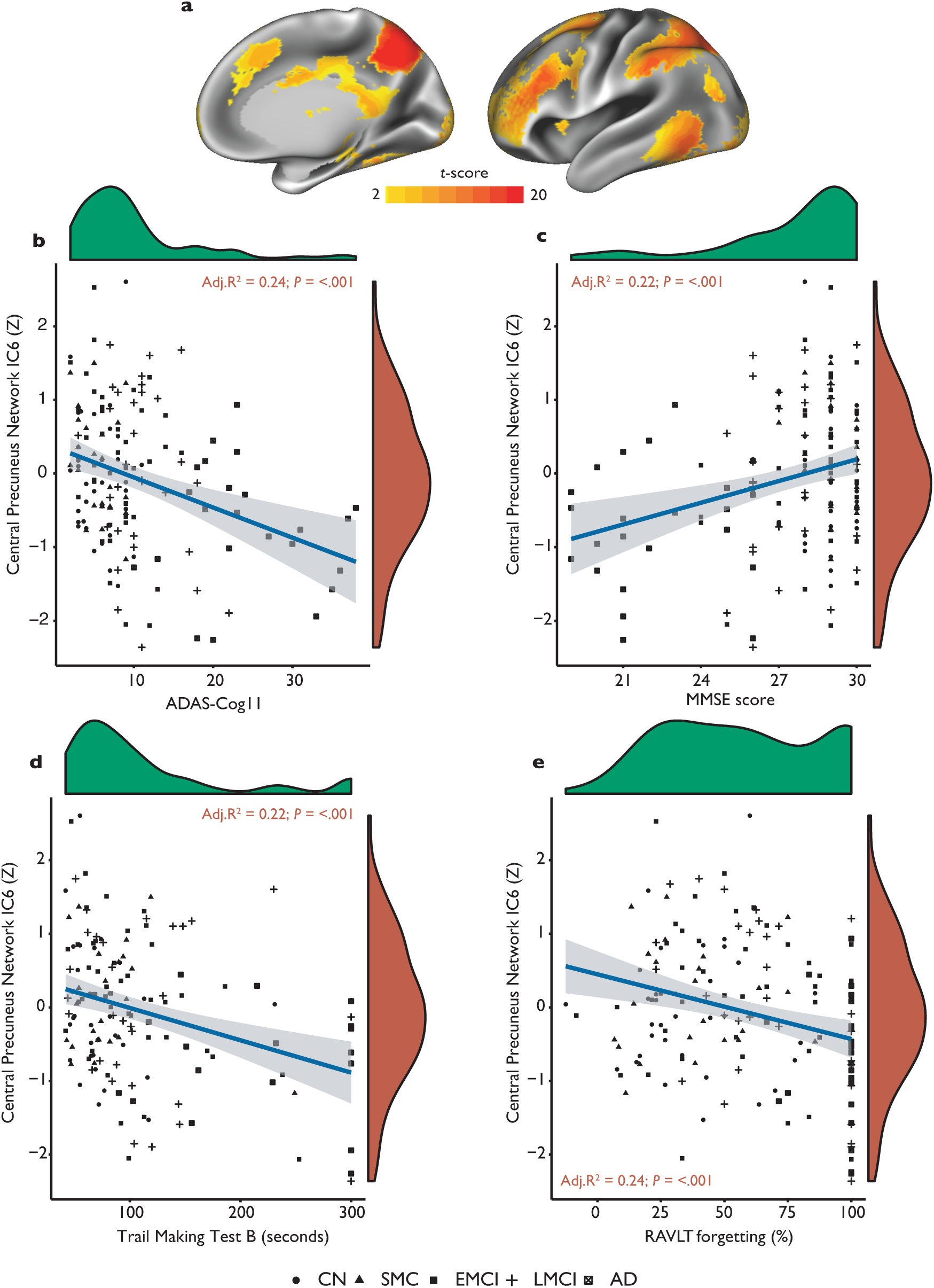
Functional connectivity of the central precuneus brain network is related to memory deficits and executive dysfunction across the Alzheimer’s disease spectrum (*N*=155). **(a)** Spatial map of the central precuneus ‘cognitive/associative’ whole-brain network is displayed on left lateral and right medial hemispheres. Functional connectivity of this network is plotted against **(b)** the 11-item Alzheimer’s Disease Assessment Scale-cognitive subscale (ADAS-cog11) scores, **(c)** Mini-Mental-State Examination (MMSE) scores, **(d)** Trail Making Test B scores and **(e)** Rey Auditory Verbal Learning Test (RAVLT) forgetting scores expressed as percentages. The density distribution as marginal plots are displayed for cognitive variables in *green* and functional connectivity *Z*-scores in *red*. Regression lines are shown in *blue* with 95% CI’s (*grey bands*). Results displayed inset are from linear regression models. Age, gender, years of education and *APOE* ε4 genotype were considered as covariates in a stepwise fashion using Akaike Information Criterion minimisation. CN = cognitively normal; SMC = subjective memory complaints; EMCI = early mild cognitive impairment; LMCI = late mild cognitive impairment; AD = Alzheimer’s disease.

## Discussion

In the current study, we used a data-driven approach to subdivide the PMC and assess how its distinct patterns of functional connectivity with the rest of the brain may be affected in AD. Previous work has also demonstrated that the PMC shows a complex functional cytoarchitecture in both human and nonhuman primates (Margulies et al., 2009; Vogt et al., 2006) and is associated with distinct functional connectivity patterns that play an active role in cognitive control (Bzdok et al., 2015; Leech et al., 2012). A unique observation from our investigation is that the functional connectivity of the PMC is selectively disrupted in AD patients. These patterns of changes were largely evident in the anterior precuneus, dorsal PCC and central precuneus subdivisions where reduced functional connectivity to large-scale brain networks was observed. In AD patients compared to CN participants, weaker functional connectivity in these subdivisions was associated with greater levels of amyloid burden and lower hippocampal volumes. Our analysis across the AD pathological spectrum (defined as ranging from CN participants and participants with SMC, through to those with different stages of MCI, and patients with a diagnosis of AD dementia) revealed that the strength of functional connectivity in the central precuneus was related to clinical disease severity, as well as specific deficits in memory and executive dysfunction. Overall these results suggest that the complex functional connectivity of the PMC is associated with multiple yet distinct large-scale brain networks that are selectively impacted in AD.

The PMC is known to be associated with distinct patterns of functional connectivity across multiple intrinsic brain networks (Leech et al., 2012; Margulies et al., 2009). In contrast to these studies, we used high resolution rsfMRI data from the HCP to create a detailed map of PMC functional connectivity patterns. ICA-based methods for decomposition are often sensitive to the number of components selected for defining the topology of intrinsic brain networks. To overcome these challenges in the arbitrary selection of ICA components, we used a data-driven approach to perform the ICA in the range of 2-20 components. This analysis was iterated 400 times to produce a global reproducibility measure that provided an estimation for extracting the ideal number of ICA components (Supplementary Methods). To the best of our knowledge, this is the first study to examine PMC functional connectivity patterns in AD using such detailed characterisations of the region.

Our findings build on existing work showing that the complex functional organisation of the PMC is related to distinct AD-related changes in functional connectivity (Vannini et al., 2013; Wu et al., 2016; Xia et al., 2014). Specifically, we found decreased functional connectivity of the dorsal PCC (IC7) in AD patients whereas the ventral PCC (IC1 and IC3) remained unaffected. The dorsal PCC is an important nexus of multiple intrinsic connectivity networks in the brain and has been proposed to modulate the global metastability of networks for efficient cognitive function (Hellyer et al., 2015). In our investigation, functional connectivity of the dorsal PCC was associated with a frontoparietal network (IC7) and a sensorimotor network (IC 10), whereas the ventral PCC (IC1) showed functional connectivity patterns that were largely consistent with the DMN. This finding is in accordance with structural and functional connectivity studies which demonstrate that the dorsal PCC exhibits selective functional connectivity patterns across a range of intrinsic networks, including frontoparietal and executive networks responsible for cognitive control (Bonnelle et al., 2011; Leech et al., 2011).

In contrast, the precuneus has been described to play a multimodal integrative role in the functional organisation of the PMC (Cavanna & Trimble, 2006; Margulies et al., 2009). Previous anatomical studies have shown that cortico-cortical connections in the precuneus have an anterior-posterior functional differentiation with largely three distinct regions (Parvizi, Van Hoesen, Buckwalter, & Damasio, 2006). A seminal study by Margulies *et al* (2009) characterised these subdivisions as an anterior zone displaying functional connectivity with sensorimotor regions, a central cognitive zone showing functional connectivity with posterior areas of the inferior parietal lobule and a posterior zone which demonstrates functional connectivity with the visual prestriate cortex and dorsolateral occipital region. This is consistent with what we observed in the functional connectivity of precuneal subdivisions of the PMC. Functional connectivity patterns of the anterior precuneus subdivision were found to be associated with a sensorimotor network (IC9) and the central precuneus (IC6) showed functional connectivity with the dorsolateral prefrontal cortex and posterior part of the inferior parietal lobule. Altered precuneus functional connectivity in AD patients has been frequently observed (Binnewijzend et al., 2012; Damoiseaux, Prater, Miller, & Greicius, 2012; Wang et al., 2006). The precuneus is also believed to play an important role in episodic memory retrieval and visual-spatial memory (S. Zhang & Li, 2012) and its involvement in memory and visual-spatial deficits in AD has been demonstrated in previous fMRI studies (Karas et al., 2007; Sperling et al., 2010). Functional connectivity differences in the precuneus have also been reported in CN participants with elevated Pittsburgh Compound B – a measure of fibrillar amyloid burden detected using PET imaging (Sheline et al., 2010).

While causative mechanisms between amyloid burden and large-scale network failures remain unclear, further examination of this association is warranted. The high degree of spatial overlap in amyloid deposition with brain regions involved in the DMN is well described in the AD literature (Buckner et al., 2005). As a result, a large number of rsfMRI studies in AD have largely focused on the relationship between amyloid burden and functional disruptions of the DMN (Hedden et al., 2009; Sheline et al., 2010). In the preclinical stages of AD, studies have also demonstrated DMN functional connectivity changes in CN participants with high levels of amyloid burden (Mormino et al., 2011). To test whether similar amyloid associations could be observed for PMC functional connectivity, we examined this relationship in AD patients and CN participants. We performed these associations for the dorsal PCC, the anterior precuneus and the central precuneus subdivisions that exhibited decreased functional connectivity in AD patients. Our findings demonstrated that weaker functional connectivity of these subdivisions was associated with greater levels of amyloid burden. Additionally, we also showed that the functional connectivity of these subdivisions was associated with lower hippocampal volumes in AD patients. This may be explained by previous work showing that functional disruptions in networks sharing a strong coherence with the hippocampus could lead to subsequent neurodegeneration in the region via a proposed ‘degenerative feedback loop’ (Damoiseaux et al., 2012; Jones et al., 2016).

Functional connectivity patterns of the PMC have only been examined in a small number of rsfMRI studies in individuals meeting MCI criteria (Petrella, Prince, Wang, Hellegers, & Doraiswamy, 2007). Moreover, no such studies have examined the progression of PMC functional connectivity patterns, ranging from CN stages through to the intermediate stages of AD development, and finally full-blown AD dementia. The functional connectivity patterns of the central precuneus subdivision, which was widely disrupted in AD patients, was used to characterise the progressive functional changes across the AD pathological spectrum. Previous evidence has also implicated the involvement of the central precuneus in higher-order cognitive and executive processing, such as the monitoring of information in working memory and action planning (Petrides, 2005). In the present study we also found a direct association between functional connectivity reductions in the central precuneus and impairments in memory and executive functioning, consistent with prior findings (Cavanna & Trimble, 2006). Furthermore, connectivity decreases were also found to be related to clinical measures of disease progression.

Recent work has shown that system-wide network dysfunctions in AD may play an aetiological role in disease pathology (Jacobs et al., 2018). One conceptualisation of such network dysfunctions proposes that a cascading network decline originates in the most highly integrated posterior brain regions (posterior DMN) and progresses via aberrant and supposedly compensatory connectivity increases of anterior DMN subsystems that trigger downstream disease pathology (Jones et al., 2016). We also demonstrate functional disruptions originating in the PMC and further highlight the vulnerability of intrinsic brain networks other than the DMN in relation to the AD clinical phenotype. However, it has been suggested that early in the disease, convergent functional hubs compensate for declines in functional connectivity by increasing cortical activity to support adjacent brain systems that may have succumbed to increased information processing loads (Schultz et al., 2017). Since our study questions were designed to characterise the functional connectivity patterns of large-scale brain networks associated with the PMC, we did not explore how connectivity patterns between nodes within this system could influence specific intra-network dysfunctions. In future work, we will address the opportunity to examine the functional connectivity of the PMC with specific nodes such as the ventromedial prefrontal cortex – a region whose increased connectivity patterns have recently been associated with higher cognitive resilience in AD (Franzmeier et al., 2018).

Some potential limitations of the current study should however be taken into consideration. In the ADNI dataset, rsfMRI scans are known to contain ‘penciling’ artefacts in part of the left lateral frontal lobe and have been observed to decrease functional connectivity in that region (Jones et al., 2016). Since we were uncertain as to how this may affect our analysis, we took all necessary precautions to avoid this region when comparing functional connectivity patterns of the PMC. We would also argue that functional parcellations of the PMC in the HCP dataset ensured that networks for the ADNI dataset were independently derived and remained topologically consistent with original network observations from HCP. Inherent limitations in fMRI scans obtained from the ADNI dataset could also be explained by the multi-centre design of the study. Although considerable efforts in ADNI were taken to harmonize acquisition protocols across different study sites, we cannot exclude the possibility that some differences in acquisition may have remained. We also only included cross-sectional data for assessing PMC functional connectivity patterns rather than using longitudinal measurements of connectivity. That being said, we have highlighted the functional importance of the PMC in AD using a range of extensively phenotyped clinical and cognitive metadata from the ADNI study.

In conclusion, our findings provide new evidence of widespread network disruptions in functionally distinct PMC subdivisions that are selectively impacted in AD. Abnormalities in the functional connectivity of the central precuneus, widely considered a cognitive processing hub, are associated with specific memory deficits and executive dysfunction across the AD pathological spectrum. Future studies should address how functional connectivity in these selectively vulnerable PMC subdivisions may potentially serve as promising future biomarkers of AD progression.

## Acknowledgements

We would like to thank Dr Andrew Simmons for his helpful discussions and advice regarding the interpretation of results. We also thank and are grateful to Chris Webb and Rodney Lewis for their work and management of the data processing clusters used for the analysis of all data in this study.

## Data Availability Statement

The data that support the findings of this study are available from the corresponding author upon reasonable request.

