## Supplementary Material for "The heterogeneous functional architecture of the posteromedial cortex is associated with selective functional connectivity differences in Alzheimer’s disease"

### Reproducibility analysis of group ICA in the HCP cohort

The mICA toolbox (version 1.17) (Moher Alsady, Blessing, & Beissner, 2016) was used to assess the ICA procedure's trade-off between granularity and noise. To determine the ideal number of dimensionalities to fractionate the posteromedial cortex signal in the HCP dataset, we performed a reproducibility analysis for a dimensionality range between 2-20 ICA components. The analysis was performed using split-half sampling which randomly splits the entire 100 HCP datasets into two groups within each sample and performs *N* repetitions across the samples. Split-half sampling was set to 20 repetitions in the dimensionality range of 2-20 components (2...4…6…8…20 = 10 ICA analyses per sample). For every repetition, MELODIC ICA analysis is performed separately on both split-half groups in each sample. Therefore, we computed a total of 10 × 2 = 20 ICA's per sample repetition and 20 × 20 repetitions = 400 ICA analyses overall.

A cross correlation matrix between the ICA spatial maps was then calculated using Pearson's coefficient. Inter-group matching of all ICA maps was performed using the Kuhn-Munkres algorithm with the Munkres package in Python 2.7 (for further details see *Moher Alsady et al., 2016*). Mean correlation of all matched IC pairs was averaged over 20 repetitions and then used as the global reproducibility measure. Confidence intervals (95%) were also generated to assess the stability of the ICA repetitions in the reproducibility curve. The final number of ICA components used for fractionating the posteromedial cortex signal were derived from the global maximum of the reproducibility curve (**Supplementary Fig. 1**).

### Alzheimer's Disease Neuroimaging Initiative (ADNI) study data

#### Image Acquisition

All subjects were scanned using a 3T Philips system. Task-free fMRI scans were obtained using a single shot gradient echo-planar imaging sequence (EPI) with full brain coverage including the whole cerebellum. The following parameters were used: 140 functional volumes, repetition/echo time of 3000/30ms, flip angle of 80°, 48 axial slices, and a 64 x 64 in-plane acquisition matrix reconstructed to produce slice thickness of 3.3mm and an isotropic 3.3mm voxel size. Further information regarding slice ordering, quality control criteria, and other ADNI protocols is provided on the www.adni.loni.usc.edu/ website.

#### Structural preprocessing

Preprocessing was carried out using a combination of the FMRIB Software Library (FSL) (Smith et al., 2004), ANTs (Avants et al., 2011) and FreeSurfer (version 6.0) (Bruce Fischl, 2012) (http://surfer.nmr.mgh.harvard.edu/). T1-weighted MRI images were acquired using the ADNI Philips 3D Magnetization Prepared Rapid Gradient Echo (MPRAGE) sequence, with 1x1mm in-plane resolution and 1.2mm slice thickness. A repetition time of ~7/3 ms was used. The Brain Extraction Tool (BET) (Smith, 2002) was used to remove non-brain tissue from the image followed by the FMRIB Automated Segmentation Tool (FAST) (Zhang, Brady, & Smith, 2001) which estimates partial volume maps of grey matter, white matter and CSF. Grey matter images for each subject were used to calculate whole-brain volume (WBV). FreeSurfer's *recon-all* pipeline was used to generate measures of hippocampal and intracranial volume. Further technical details of FreeSurfer's cortical reconstruction and segmentation procedure have been described in previous publications (Dale, Fischl, & Sereno, 1999; B Fischl et al., 2002; B Fischl & Dale, 2000; Reuter, Rosas, & Fischl, 2010; Ségonne et al., 2004). A custom study-specific template was constructed using ANTs advanced normalisation tools (Avants et al., 2011) for more accurate registration of structural and functional images (Klein et al., 2009). This template was constructed to the dimensions of Montreal Neurological Institute (MNI) space to allow for whole-brain maps of functional connectivity to be compared with the same whole-brain maps derived from the HCP data.

### Functional preprocessing

Preprocessing of rsfMRI data was carried out using FSL. The first 10 volumes were discarded for steady-state magnetization and to avoid including volumes contaminated with scanner artefact. Standard preprocessing steps included motion correction, non-brain tissues removal, spatial smoothing with a 6mm full width at half maximum Gaussian (FWHM) kernel, and high-pass temporal filtering with a cut-off frequency of 0.01 Hz. Subsequently, single-subject spatial ICA with automatic dimensionality estimation was performed using MELODIC (Beckmann, DeLuca, Devlin, & Smith, 2005) and FMRIB's ICA-based Xnoiseifier (Salimi-Khorshidi et al., 2014) for the removal of artefactual components. FIX was manually trained on 25 rsfMRI datasets (Mean Age: 72.7 ± 6.1 years, Male/Female = 14/11) and consisted of 5 datasets randomly selected from each group (CN-SMC-EMCI-LMCI-AD). The accuracy of FIX classifications were assessed manually (Griffanti et al., 2014). Motion artefacts were also assessed by examining fMRI volumes that may potentially be corrupted by motion using parameters of Framewise Displacement (FD) and DVARS which measures rate of change in BOLD signal across the entire brain at each timepoint of the data (Power, Barnes, Snyder, Schlaggar, & Petersen, 2012). Our criteria for excessive motion included > 0.50mm in more than 15% of volumes and/or an FD > 3.0mm across any of the volumes (De Simoni et al., 2018;

Manza, Zhang, Li, & Leung, 2016). In total, 17 subjects were removed using these criteria leaving a final ADNI fMRI subset of 155 participants.

After preprocessing, each single-subject 4D dataset was first aligned to the subject's high-resolution T1-weighted image using FMRIB's Linear Image Registration Tool (FLIRT) (Jenkinson, Bannister, Brady, & Smith, 2002; Jenkinson & Smith, 2001) which was enhanced with brain-boundary registration (Greve & Fischl, 2009). These images were then registered to the custom study-specific template resampled to MNI152 standard space ($2 \times 2 \times 2mm^3$) using FMRIB's Nonlinear Image Registration Tool (FNIRT).

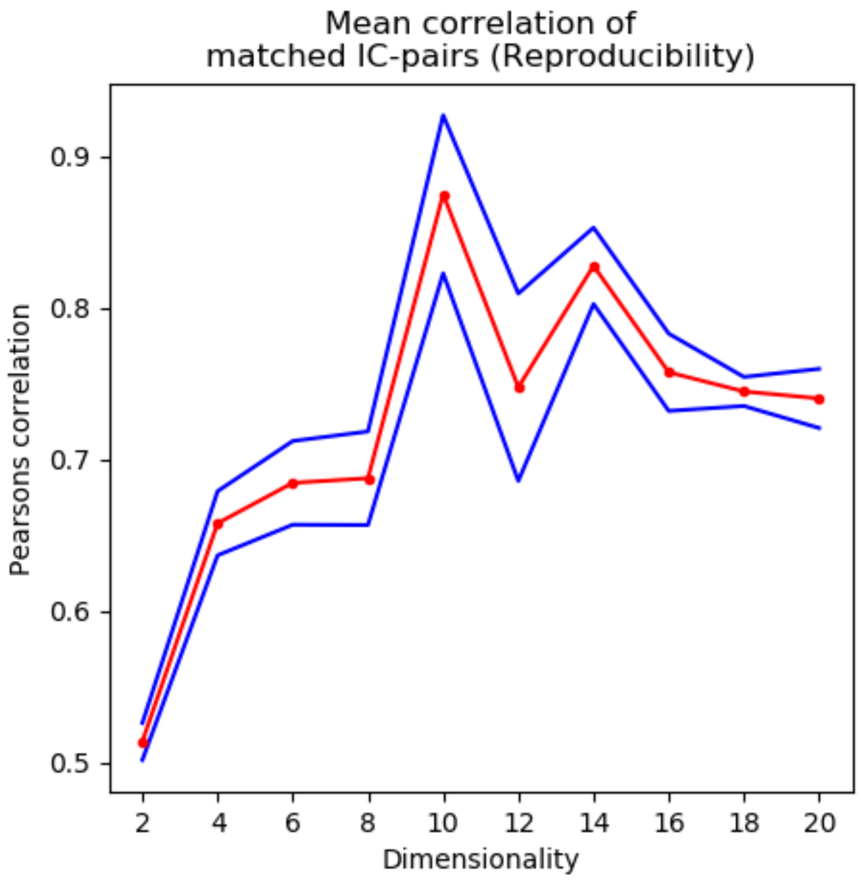


**Supplementary figure 1 reproducibility analysis of the ideal number of ICA components for subdividing the posteromedial cortex.** The ideal number of ICA components to use for fractionating the PMC were decided from the dimensionality-reproducibility curve (red = mean, blue = 95% CI's), where the 10 component ICA was the global maximum. Therefore, the 10 component ICA analysis was chosen as the ideal number of components to subdivide the PMC into its distinct subdivisions.
